# Continuous evolution of a halogenase enzyme with improved solubility and activity for sustainable bioproduction

**DOI:** 10.1101/2025.10.08.681035

**Authors:** Andre Arashiro Pulschen, Justin Booth, Ari Satanowski, Christelle Soudy, Joaquin Caro-Astorga, Osaid Ather, Namita Patel, Ali Alidoust, Samir Aoudjane, Lily Nematollahi, Erika DeBenedictis

## Abstract

Halogenation enhances the stability and function of pharmaceuticals, biomaterials, and industrial compounds. However, chemical halogenation lacks stereoselectivity and requires the use of toxic or expensive chemicals. Although enzymatic halogenation can improve selectivity and reduce environmental impact, current halogenases are inefficient and insoluble, leading to low yields that limit their applications. Here, we develop RebH_Evo4_, a soluble and highly active tryptophan halogenase, containing 12 mutations that confer 37-fold and 44-fold increases in 7-chloro and 7-bromotryptophan production respectively, *in vivo.* To create RebH_Evo4_, we devised an aminoacyl tRNA synthetase based halogenase biosensor and conducted over 500 hours of phage-assisted continuous evolution (PACE). Use of RebH_Evo4_ in a bioreactor resulted in the production of 2.7 g/L of halogenated tryptophan. When coupled with a downstream enzyme, RebH_Evo4_ allowed 36-fold increased yields of halogenated tryptamines compared to the wild-type enzyme. Additionally, RebH_Evo4_ enabled efficient production of genetically encoded antimicrobial halogenated peptides. The efficient, site-specific halogenation by our evolved halogenase will accelerate sustainable biomanufacturing of halogenated drugs.

## Introduction

Halogenated compounds are central to modern medicine, with about 25% of approved drugs and over 80% of agrochemicals incorporating halogens to enhance bioavailability and stability^1,2,3^. While chemical halogenation is a well-established method, it often requires harsh conditions involving toxic and corrosive reagents^4,5^ and can lack regio- and enantioselective control. Enzymatic halogenation is emerging as a powerful, stereoselective, and environmentally sustainable alternative^5^. Tryptophan halogenases such as RebH (which can natively halogenate the tryptophan indole group at position 7 with both chloride and bromide) are among the best-studied biocatalysts for halogenation. These enzymes rely on a two-step catalysis, first oxidising a flavin cofactor to drive production of an intermediate hypohalous acid (HOX) species, then ensuring precise electrophilic halogenation at the indole ring of tryptophan and related compounds^6^.

Tryptophan and its derivatives are key precursors of several molecules of interest, including those with applications in the food industry, pigments^7,8^, agriculture, and medicine^9,10,11,12^. Halogenation of Tryptophan or indole is necessary for the production of important molecules such as the pigment tyrian purple^7^, and the antifungal pyrrolnitrin^11^. Additionally, Tryptophan halogenation has been explored for the diversification of natural products, which can improve stability, pharmacokinetics, and activity^5^. It has also been explored for the diversification of other active molecules, like violacein^8^ and plant monoterpenes, such as alstonines and serpentines^13^, carbolines and pro-drugs halogenated kynurenines^12^. Selective incorporation of halogenated tryptophan can also improve protein and peptide drugs. For example, a recently discovered halogenated variant of the antibiotic Darobactin A is more active than its non-halogenated compound^14^. Krisynomycin and Nisin are additional examples of molecules in which their activity and stability is improved due to halogenation^15,16^

To date, however, there has been limited success in using flavin-dependent halogenases to produce substantial amounts of halogenated tryptophan compounds *in vivo.* This is due to poor solubility, low enzymatic activity, and temperature sensitivity^17^, leading to low fermentation yields^17^ and slow reaction rates^7,12^. Although efforts have been made to engineer halogenases with expanded substrate scope^18,19^, enhanced thermotolerance in *in vitro* conditions^20^, and decreased leakiness of intermediate HOX^21^, it remains a challenge to obtain highly active variants. As a consequence, despite their limitations, wild-type sequences continue to comprise the majority of enzymes used for bioproduction and fermentation^7,8,12,13,22^. Notably, there are no described variants with improved solubility; instead, fusion with more soluble proteins, such as maltose-binding protein and flavin reductases, and co-expression of chaperones are used in an attempt to overcome this^7,23,24^.

In this study, we sought to overcome these challenges by applying continuous directed evolution to the flavin-dependent tryptophan halogenase RebH, To establish a readout for halogenation activity, we developed a biosensor based on the engineered aminoacyl-tRNA synthetase ChPheRS-4^25^, coupling the biosynthesis of the halogenated non-canonical amino acid (ncAA) 7-chlorotryptophan (7-Cl-Trp) to expression of the sfGFP gene. We then adapted our biosensor to link the activity of RebH to phage propagation, allowing us to conduct over 500 hours of Phage-Assisted Continuous Evolution (PACE). Our final evolved variant, RebH_Evo4_, contains 12 mutations and exhibits improved solubility and activity compared to the wild-type enzyme. When coupled with the downstream enzyme RgnTDC, it allowed the efficient production of halogenated tryptamines. Additionally, we used RebH_Evo4_ for bioproduction of halogenated antimicrobial peptides, demonstrating its use for the efficient manufacture of proteins with site-specific halogenation in *E. coli*.

## Results

### Aminoacyl-tRNA synthetase biosensor to detect halogenation *in vivo*

We envisioned that an aminoacyl-tRNA synthetase (aaRS) that is selective for a halogenated ncAA could function as a biosensor, enabling amber stop codon (TAG) suppression only when a halogenated amino acid is added to the media, or halogenation occurs *in vivo*. We designed a genetic circuit to utilize the engineered chimeric aaRS ChPheRS-4, which has previously been reported to incorporate 7-Cl-Trp into proteins at TAG codons^25^ (Figure 1A). By incorporating a TAG amber stop codon within the gene encoding sfGFP, using a GFP-151-TAG cassette, amber suppression using the halogenated ncAA allows full sfGFP translation and thus fluorescence, which can be used as an indirect measure of tryptophan halogenation. In our hands, using a Δ*tnaA* (tryptophanase deletion) strain to prevent tryptophan degradation, ChPheRS-4 demonstrated weak amber suppression, as measured by GFP signal (Supp. Figure 1A). To improve the biosensor, we searched the literature for variants of the aaRS-tRNA pair that have been shown to improve amber suppression on related ncAAs and identified two possible variants: the aaRS mutation S333C and the tRNA mutation 3C11^26^. Testing these variants confirmed improvements in amber suppression and thus 7-Cl-Trp incorporation into sfGFP (Supp. Figure 1A and 1B). As media composition is known to strongly impact the performance of amber suppression circuits, we compared amber suppression in five different types of media (Figure 1B). The largest fold-change occurred using

**Figure 1.**
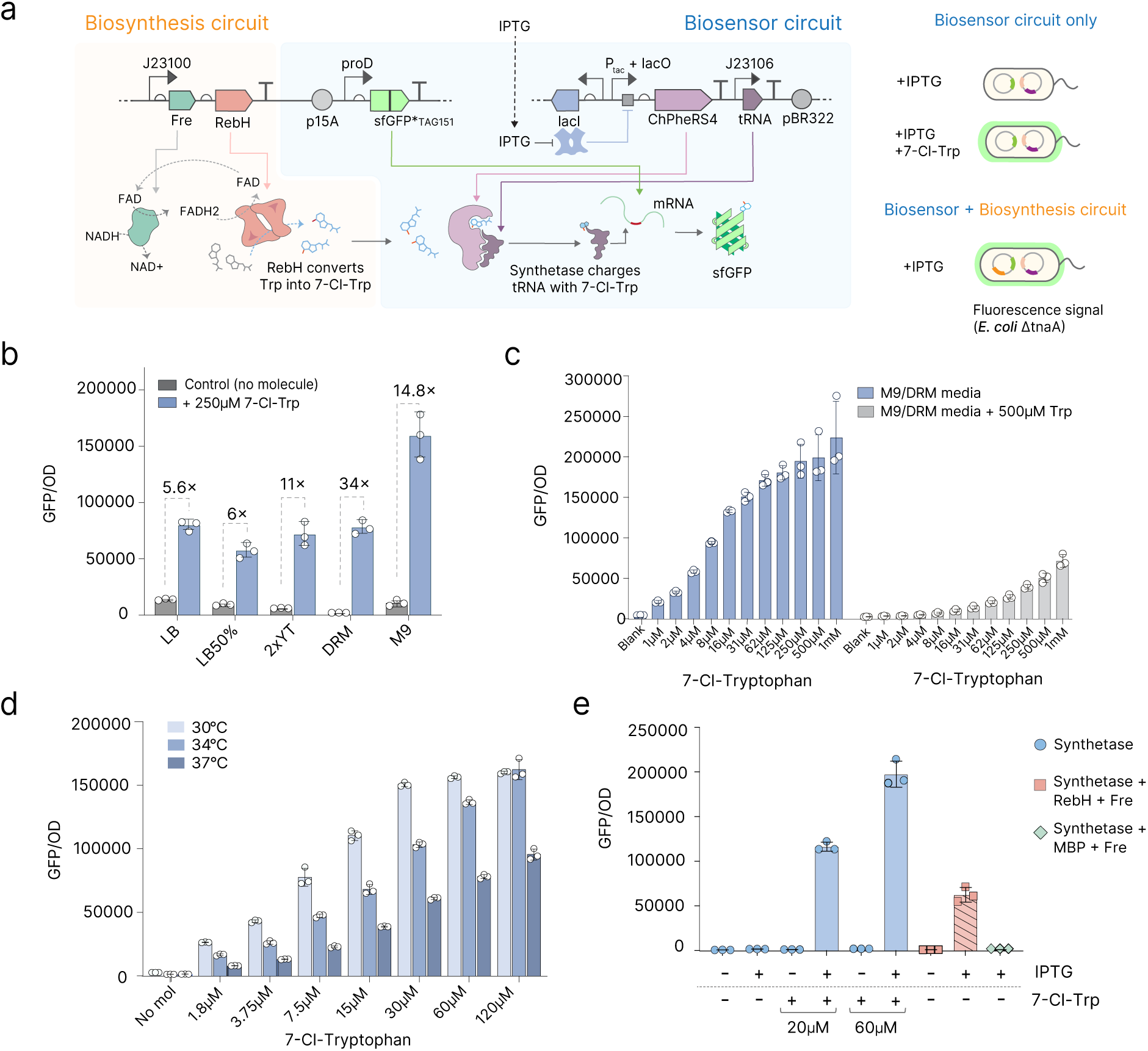
Generation of an AARS-biosensor and biosynthesis circuit for halogenation. 1a. Schematic of our aaRS-based biosensor for 7-Cl-Trp, and a circuit for biosynthesis of 7-Cl-Trp using the flavin-dependent halogenase RebH that results in amber codon suppression, incorporation of 7-Cl-Trp, and thus full-length expression of sfGFP. The biosensor can be tested independently or combined with the biosynthesis pathway in the same *E. coli* cell. 1b. Measurement of biosensor readout in different media backgrounds. The graph depicts GFP normalized by OD_600_. Error bars show mean and SD. 1c. Measurement of the biosensor readout in a 10:90 DRM/M9 media background, with or without supplementation with canonical tryptophan. N=3 biological replicates. 1d. Measurement of the biosensor readout at 30°C, 34°C, and 37°C, with different concentrations of 7-Cl-Trp.Error bars show mean and SD. 1e. Measurements of the biosensor and biosynthesis circuit within the same cell. Bars indicate the use of the synthetase (ChPheRS-4), Halogenase (RebH), Flavin reductase (Fre), and Maltose Binding Protein (MBP). Error bars show mean and SD.

DRM media, which was originally formulated for PACE experiments^27^, while the largest absorbance-normalized GFP signal was observed when using M9. We chose to combine both types of media in a 1:9 DRM/M9 ratio, which allowed strong GFP signal, with very low leakiness (Figure 1C). The resulting biosensor enables robust detection of 7-Cl-Trp at concentrations as low as 1 µM (9-fold increased signal over background) and with the operational range reaching as high as 125 µM to 250 µM (95-fold higher signal over background), at 30°C (Figure 1C).

We observed that GFP signal was substantially reduced when the media was supplemented with canonical L-tryptophan in addition to 7-Cl-Trp (Figure 1C), indicating that non-halogenated tryptophan acts as an inhibitor of the ChPheRS-4 synthetase and, in effect. Additionally, sfGFP signal was abolished when 7-Cl-Trp was replaced with tryptophan, indicating that the aaRS does not promiscuously charge tRNAs with a non-halogenated substrate. These observations indicate that negative selection against incorporation of canonical tryptophan is not necessary for this aaRS, and that the stringency of our biosensor can be adjusted with the simple addition of canonical tryptophan to the media.

We further characterized our biosensor by measuring its activity at different temperatures. In previous work, amber suppression has been measured at 30°C^25^. In our hands, we found that the efficiency of amber suppression is reduced at 34°C and 37°C when compared to 30°C (Figure 1D). However, no GFP signal was detected in any of the tested temperatures in the absence of the ncAA, confirming that overall synthetase fidelity is maintained throughout this temperature range (Figure 1D).

Moving on from exogenous addition of 7-Cl-Trp, we next sought to confirm that our biosensor could be used to measure the activity of halogenase enzymes expressed *in vivo*. Since all known tryptophan halogenases rely on an available pool of reduced flavin cofactors, co-expression with a flavin reductase has been shown to improve *in vivo* yields of halogenated tryptophan^7,28^. We co-expressed the flavin-dependent halogenase RebH (*Lechevalieria aerocolonigenes,* Uniprot Q8KHZ8) with a flavin reductase (Uniprot P0AEN1) in *E. coli* (Figure 1A), and found that expression of the halogenase inside the cell results in a GFP signal that is similar in magnitude to direct addition of 7-Cl-Trp to the media, suggesting that the halogenase was actively producing this halogenated ncAA (Figure 1E). To further confirm that the GFP production is due to the presence of RebH and not the flavin reductase, we tested a construct replacing RebH with the innocuous maltose binding protein (MBP), and confirmed that no GFP production was observed (Figure 1E). Together, these data demonstrate that the biosensor enables robust measurement of halogenase activity and that 7-Cl-Trp biosynthesis occurs *in vivo*.

### Halogenation-dependent phage propagation

Next, we sought to adapt the GFP readout of our biosensor to an M13 phage readout, a prerequisite for conducting halogenase evolution using PACE. PACE^29^ is a continuous evolution method in which the protein to be evolved is encoded in place of the pIII-encoding gene in an M13 bacteriophage. The resulting loss of pIII protein production prevents replication of the phage unless pIII is produced by a circuit encoded in the *E. coli* cell it infects. Since circuit pIII output is constructed to be dependent on the activity of the protein of interest, phage propagation is linked to the function of the protein being evolved. When propagated against a constantly-replaced pool of host cells under high mutagenesis, PACE allows for rapid exploration of the sequence-function landscape and selection of activity-improving mutations^27^.

For our circuit, we expressed RebH on the phage under the strong constitutive proK promoter, while the aaRS-tRNA pair and flavin reductase are encoded by accessory plasmids. The sfGFP_*TAG151_ from our previous circuit was replaced with pIII_*TAG29_, which has previously been used to tie amber suppression to M13 phage propagation^30^. Upon infection of *E. coli* host cells (K-12 Δ*tnaA* cells + F plasmid), RebH is expressed from the phage and induces amber suppression by production and translational incorporation of 7-Cl-Trp, thereby allowing production of pIII and phage propagation (Figure 2A). We tested this design using phage plaquing and confirmed that RebH-encoding phage can successfully propagate without the addition of ncAA to the media, whereas a negative control empty phage (lacking RebH) could only propagate when 7-Cl-Trp was added to the media (Figure 2B). Since RebH is known to be a highly temperature-sensitive enzyme, we next probed the temperature-dependence of this system. At 34°C, phage propagation was very limited, but still detectable for the phage encoding RebH, whereas an empty phage suffered from strong de-enrichment (Figure 2C). Finally, phage propagation at 37°C was only observed upon addition of 7-Cl-Trp to the media (Figure 2C). These data indicate that 34°C is a viable starting temperature for evolution, at which all components of the system are sufficiently active, with a maximum possible phage enrichment of three-log-fold phage enrichment upon sufficient 7-Cl-Trp presence (Figure 2C).

**Figure 2.**
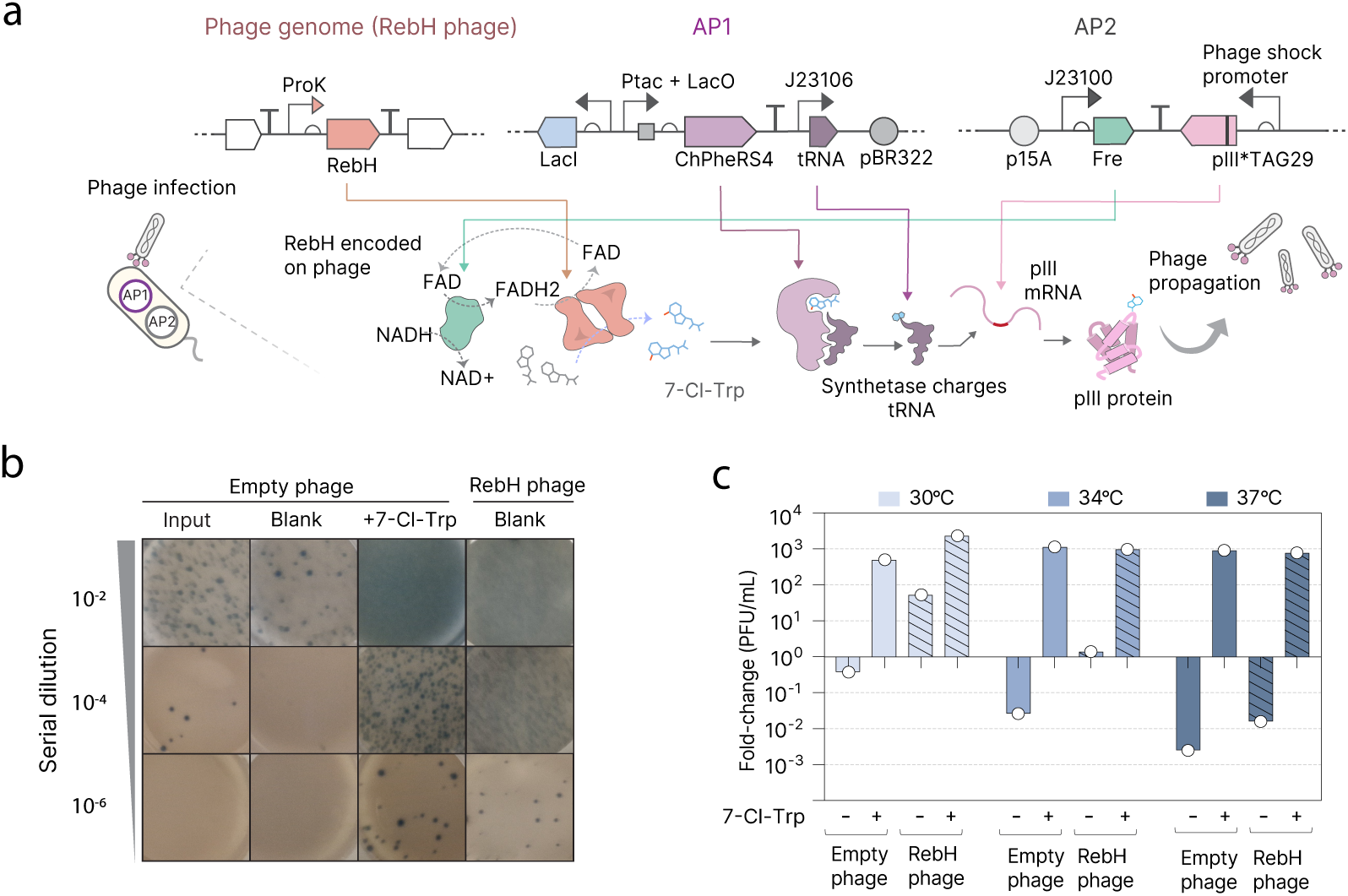
Halogenase M13 phage-based gene circuit. 2a. M13 phage-based halogenase circuit. RebH gene is encoded on the phage; Accessory Plasmid (AP) 1 produces aaRS and tRNA, AP2 produces Fre and pIII mRNA containing a single amber strop codon at position 29. Only 7-Cl-Trp production allows amber suppression of the pIII_*TAG29_ and thus phage propagation. 2b. Plaque results of an overnight phage enrichment at 30°C performed in liquid DRM:M9 media in the presence of phage bearing RebH or no gene, and with either no small molecule added or direct addition of 7-Cl-Trp. 2c. Overnight phage enrichment (in DRM:M9 media) across different temperatures and in the presence or absence of phage-encoded RebH or supplementation with 7-Cl-Trp (200µM).

### Phage-assisted Continuous Evolution of RebH

We evolved RebH using PACE for a total of 560 hours of continuous flow divided over three phases (Figure 3A). In the first phase, we evolved WT RebH with continuous *in vivo* mutagenesis induced by the mutagenesis plasmid MP6^31^ for 240 hours (Figure 3B), gradually increasing the temperature from 34°C to 37°C to push the selection to more thermostable variants. Sequencing 24 clonal phage from the final timepoint, we observed 2 co-occurring mutations that fixed in the population: V256I and T385I (Supp. Figure 2A). We also sequenced clonal phage from an intermediate sample taken at 160 hours and observed additional mutations (M430L, T348A, L188F, and L380F) that did not reach fixation. We tested combinations of all six mutations using our GFP biosensor circuit (Supp. Figure 2B) and obtained a further improved variant with 4 mutations (256I, 348A, 385I, 430L), which we called RebH_Evo1_, that demonstrated improved signal (Figure 3C, 3D).

**Figure 3.**
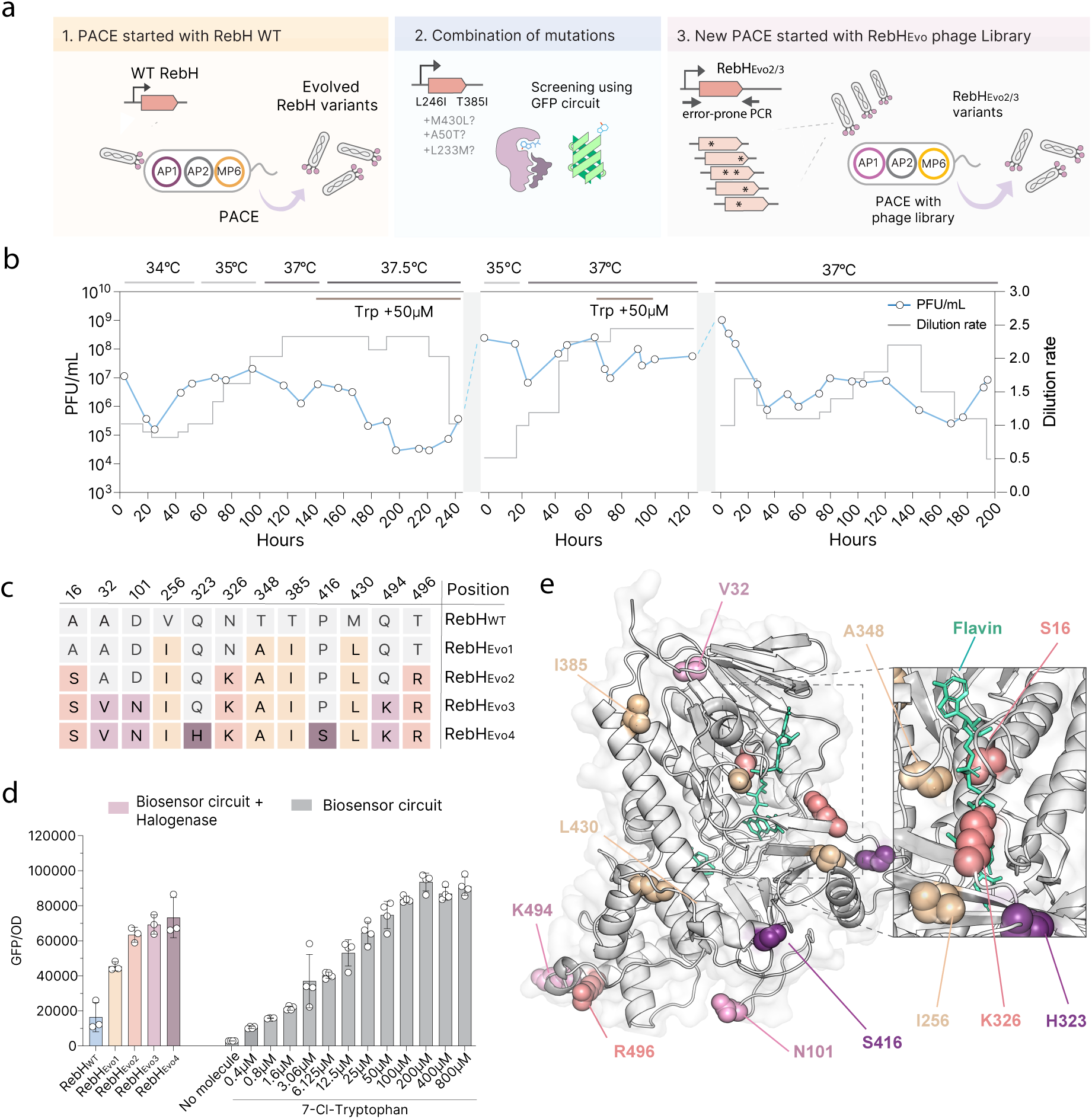
Phage-assisted continuous evolution of RebH halogenase. 3a. Schematics of the evolution campaign. 3b. Phage titres (measured by qPCR), dilution rates, and temperatures during the different PACE runs. 3c. Mutations across RebH during evolution campaigns. 3d. GFP/OD measurement of evolved RebH variants obtained during PACE campaigns, at 37°C, in DRM:M9 media (10:90). Measurements are displayed side-by-side with direct addition of 7-Cl-Trp to the media for comparison. Error bars show mean and SD. 3e. Predicted structure of RebH_Evo4_, generated by Alphafold2, with mutations highlighted.

The success of combining different mutations observed from the first phase of PACE led us to use a similar approach in our second and third phases of continuous evolution, where we initiated PACE with pre-diversified libraries (Supp. Figure 2C), resulting in successful generation of a larger diversity of phage with several mutations relative to initiating evolution with a clonal genotype (Supp. Figure 2A).

In our second phase, we templated an error-prone PCR library on the RebH_Evo1_ mutant and used this library to initiate continuous evolution. After 120 hours of continuous flow, we analyzed the sequence of 40 clonal phage. Among the sequences, we observed several instances in which independent phage had mutations at very close positions structurally (Figure 3E), such as D101N and G102S, and T496R and Q494K (Supp. Figure 2D). The use of pre-diversified libraries allowed for several different mutations to be selected in the second PACE round, but none were fixed in the population. We then tested a combination of the mutations found in the few most enriched phage, to identify if any of their improvements were additive (Supp. Figure 2D), and we obtained RebH_Evo2_, containing three additional mutations that arose during the second phase (7 mutations total, Figure 3C). Building on RebH_Evo2,_ we tested a combination of other mutations found in the second PACE round to obtain RebH_Evo3_, a variant containing 10 mutations relative to RebH_WT_, that achieved the highest activity of all variants tested.

Finally, in our third PACE phase, we repeated the process using RebH_Evo3_ as the template for library diversification, which was used to initiate evolution. After 200 hours of continuous flow, we obtained one fixed mutation (P416S) and a few enriched variants. After a new round of GFP screening and mutation combination (Supp. Figure 2E), we obtained our final RebH variant, RebH_Evo4_, with a 5-fold increase in sfGFP signal compared with the WT enzyme (Figure 3D and Supp. Figure 2E). However, since our sfGFP circuit response is not linear at high 7-Cl-Trp concentrations (Figure 3D), we anticipated the real increase in RebH activity to be much higher. Our final evolved RebH_Evo4_ enzyme contains 12 mutations (A16S, A32V, D101N, V256I, Q323H, N326K, T348A, T385I, P416S, M430L, Q494K, and T496R), which are spread across the protein structure (Figure 2F). Some of these mutations were observed at surface residues (Q494K, T496R, P416S, D101N, and Q323H), whereas some face the internal structure of the protein (T385I, A32V, and M430L). Interestingly, at least four mutations were observed around the flavin binding pocket (T348A, A16S, V256I, and N326K), with T348A and A16S having their side chain very close to flavin itself (Figure 2F).

### Evolved RebH_Evo4_ is more soluble and more active than the WT

We next sought to characterize our evolved halogenase RebH_Evo4_ in a whole-cell bioconversion assay using HPLC/MS. K-12 ΔtnaA cells encoding Fre and either RebH_Evo4_ or WT RebH were grown to mid-log phase, transferred to a minimal biocatalysis buffer, and allowed to catalyse overnight reactions with saturating tryptophan. RebH_Evo4_ exhibited a 37-fold increase in production of 7-Cl-Trp at 37°C compared to the WT enzyme (Figure 4A and 4B, Supp. Figure 3), demonstrating a substantial gain in overall activity. We also tested the activity of our enzyme at 37°C versus the WT at 30°C, which is the optimal temperature for WT RebH expression in *E. coli* due to its poor solubility^12,7^. Reassuringly, the activity of our evolved enzyme at 37°C still outperformed the WT enzyme at 30°C by 12-fold (Figure 4A and 4B and Supp. Figure 3).

**Figure 4.**
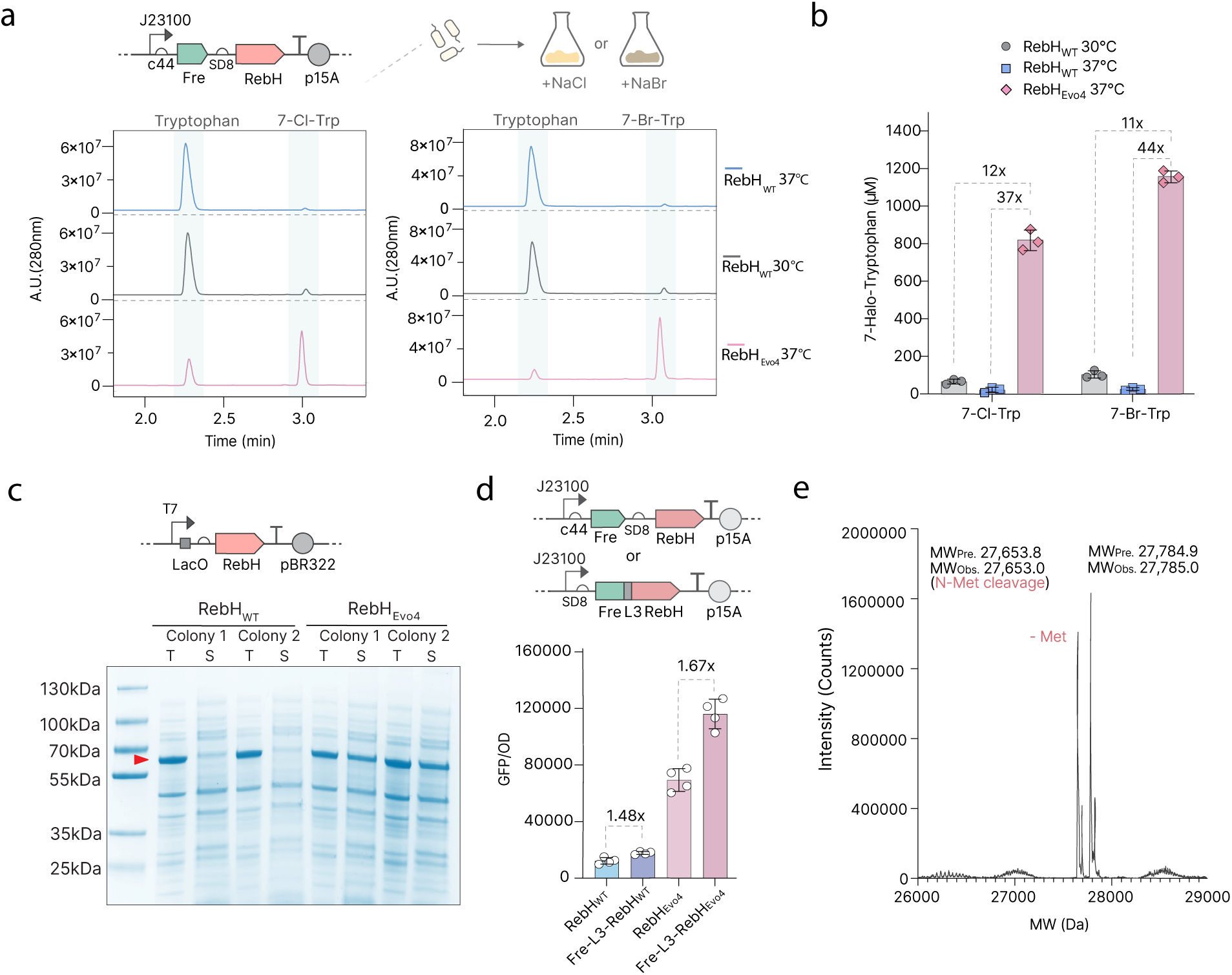
Characterization of evolved halogenase RebH_Evo4_. 4a. HPLC of whole-cell expressing RebH_WT_ or RebH_Evo4_ at 30°C or 37°C, for the production of 7-Cl-Trp (left) or 7-Br-Trp (right). 4b. HPLC quantifications of 7-Cl-Trp and 7-Br-Trp production by whole-cell reaction (starting OD600_nm_ of 5). Error bars show mean and SD. 4c. Comparison of solubility of RebH_WT_ and RebH_Evo4_ as measured by PAGE gel, using BL21 (DE3) strains and T7 promoter. T = Total fraction, S= Soluble fraction. 4d. Comparison between Fre-L3 fused and non-fused RebH variants in the GFP biosynthesis and biosensor system expressed in the K-12 strain of *E.coli*. Error bars show mean and SD. 4e. Whole-protein mass spectrometry of sfGFP produced using Fre-L3-RebH_Evo4_. The correct expected mass for 7-Cl-Trp incorporation at position 151. MW_Pre._: Predicted molecular weight. MW_Obs._: Observed molecular weight.

Next, we measured RebH_Evo4_’s ability to halogenate using bromide instead of chloride. We replaced NaCl with NaBr in the media, and found that yields for the brominated product were also increased using RebH_Evo4_, with a 44-fold increase compared to the WT at 37°C and an 11-fold increase at 30°C (Figure 4A and B, Supp. Figure 4).

RebH is known to be an extremely insoluble protein^7^. Since we observed several surface mutations in our evolved mutant, we investigated whether PACE had affected the overall solubility of the protein. We overexpressed the enzyme variants and observed that RebH_Evo4_ is significantly more soluble than WT as measured via SDS-PAGE (Figure 4C).

The activity of tryptophan halogenases has been improved by N-terminal fusion of a flavin reductase^7^. Two mechanisms may be responsible: improved solubility or creation of a channelling effect between flavin and the halogenase, thus increasing localized concentrations of FADH_2_^24^, an essential cofactor for RebH. We tested whether N-terminal fusion of a flavin reductase can further improve the activity of RebH_Evo4_, using the previously described L3 linker^7^. Indeed, we found that WT RebH and RebH_Evo4_ are both improved by the fusion, with similar fold-change (Figure 4D; 1.5-fold for WT and 1.7-fold for RebH_Evo4_), suggesting that such improvements at 37°C are likely due to solubility-independent channeling effects.

Finally, we tested the efficiency of amber suppression of our evolved enzyme at 37°C. To improve the signal in this assay, we added a new point mutation, M490L, to the ChPheRS-4_*S333C_ synthetase, which has been reported to increase its efficiency at 37°C^26^. Running our K-12 ΔtnaA strain in a modified M9 medium, we reached 100% amber suppression in cells expressing the full biosensor-biosynthesis circuit with 1 TAG present, and 20% when 3 TAGs were used; the WT enzyme can only achieve 22% amber suppression with 1 TAG (Supp. Figure 5A). We then confirmed by mass spectrometry that the sfGFP produced had correctly incorporated 7-Cl-Trp and 7-Br-Trp (Figure 4E and Supp. Figure 5B). These results indicate that our evolved enzyme enables complete site-specific halogenation at one residue, as well as site-specific halogenation at multiple residues, neither of which was possible with the WT enzyme.

### Efficient production of halogenated biomolecules and peptides using RebH_Evo4_

Enzymatic halogenation is a promising alternative to chemical methods for inexpensive, green production of commodity chemicals. We sought to explore the utility of our evolved RebH_Evo4_ beyond its native product, 7-halotryptophan. Halogenated tryptamines are an important class of biomolecules, which are widely used as drugs for migraines and cluster headaches^33^, and have recently begun to attract attention owing to the success of serotonin receptor agonists in combating treatment-resistant depression^34^. Indeed, 7-chlorotryptamine is one of the most potent serotonin 5-HT_2A_ receptor agonists reported to date^35^. Halogenated tryptamines have also been used as precursors for the discovery and production of several active molecules derived from plants, such as alstonine^36^ and the anti-cancer precursor strictosidine^37^. Using a 2 plasmid system, we coupled our halogenase and flavin reductase with RgnTDC, a tryptophan decarboxylase, to create a metabolic pathway converting tryptophan into 7-chloro or 7-bromotryptamine^12,38^. Just like with its native product, we observed that RebH_Evo4_ dramatically increased the final yields, with a 24-fold and 36-fold increase in 7-chloro and 7-bromotryptamine production, respectively (Figure 5A), indicating that the utility of RebH_Evo4_’s improvements are not limited to its native product and may have wider applications in metabolic engineering.

**Figure 5.**
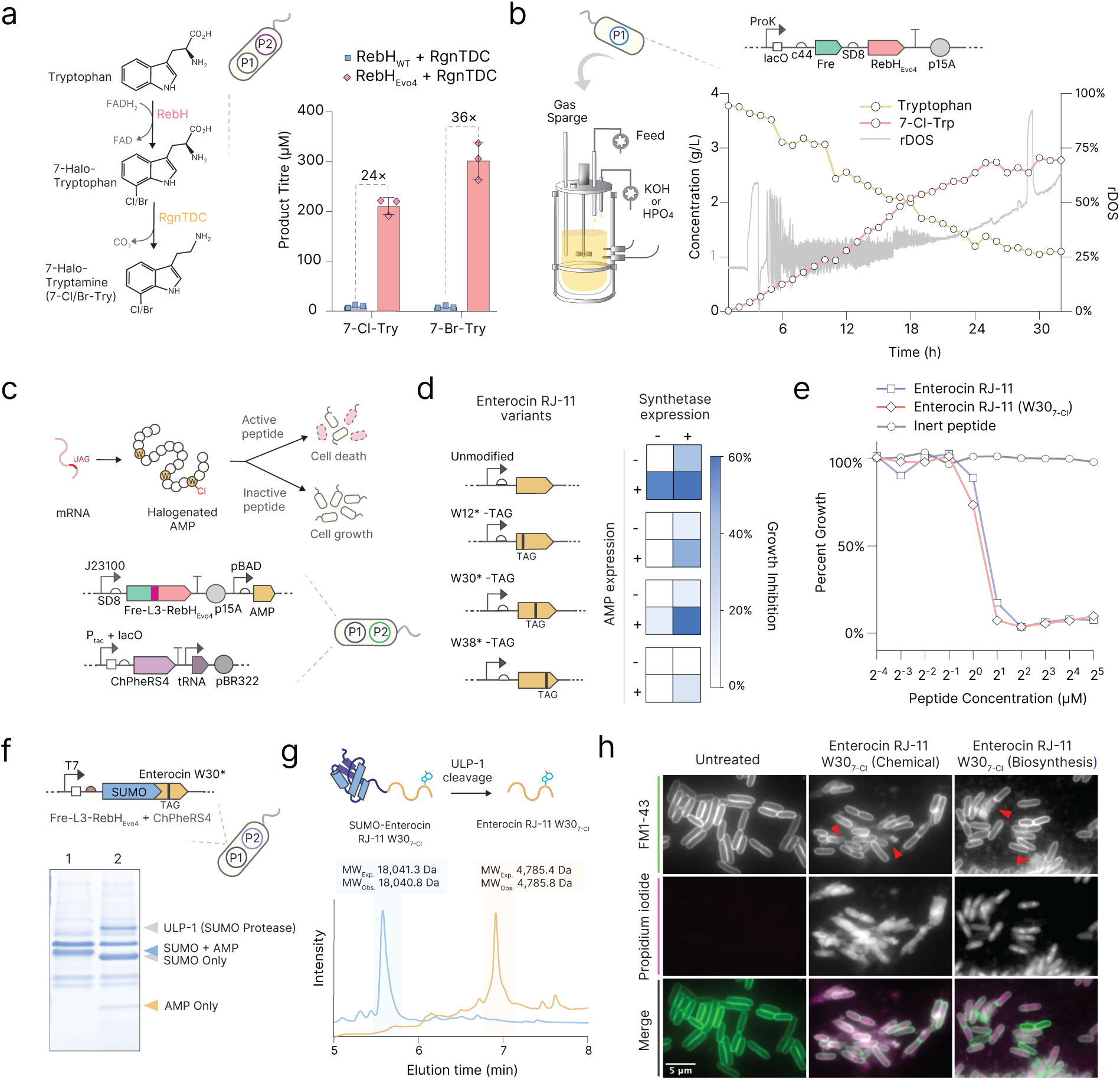
Bioproduction of halogenated biomolecules and halogenated antimicrobial peptides. 5a. Production of halogenated tryptamine using unfused RebH, Fre and RgnTDC, at 37°C. Error bars show mean and SD. 5b. Fermentative production of 7-chlorotryptophan in a fed-batch bioreactor process, at 37°C. 5c. Schematic of halogenated AMP suicide assay. Halogenated AMP variants are expressed *in vivo* and active variants identified via growth suppression 5d. Growth rate suppression data (% relative to untransformed control strain) for halogenation and truncation at all three Trp residues in the AMP Enterocin RJ-11 5e. MIC assay showing *E. coli* kill curves of commercially-synthesised WT and Trp30-Cl Enterocin RJ-11, alongside an inert control peptide 5f. SDS-PAGE analysis of halogenated peptide purification intermediates. Left lane(1) is whole IMAC eluate, right lane(2) is eluate post SUMO cleavage with ULP-1. 5g. HPLC/MS of SUMO-peptide pre and post cleavage and purification showing production of the correctly purified halogenated species. MW_Pre._: Predicted molecular weight. MW_Obs._: Observed molecular weight. 5h. Fluorescence microscopy assay showing *E. coli* cells treated with commercially chemically-synthesised Trp30-Cl Enterocin RJ-11 and purified biosynthesised product. Cells stained with FM 1-43 and propidium iodide to mark disrupted membranes.

To further explore the titres that our evolved RebH_Evo4_ was capable of producing and assess whether it would be compatible with industrial production, we scaled up our bioconversion process to a 5L fed-batch bioreactor, feeding a culture in our M9-based minimal medium using relative dissolved oxygen saturation control (rDOS-stat) with a glucose-based feed. Over the course of approximately 30 hours at 37°C, we obtained a final titre of 2.7 g/L (approx. 12 mM) 7-Cl-Trp at (Figure 5B). To our best knowledge, this represents the highest titre currently reported in the literature^7,12,17,39^.

We then decided to apply our evolved halogenase towards the production of halogenated proteins. Antimicrobial peptides (AMPs) have emerged as a promising modality for combating multi-drug resistant pathogens, however they characteristically suffer from instability and low protease resistance^40^. Tryptophan halogenation is observed widely in natural AMPs and has been demonstrated to improve these traits^14,16^. For example, the AMP Krisynomycin contains two 7-Cl-Trp residues which are crucial for its potent activity against methicillin-resistant Staphylococcus aureus (MRSA)^15,41^. However, engineering halogenation into novel AMPs is challenging since finding positions which tolerate this modification and producing successful variants at scale requires slow-turnaround custom solid phase synthesis using ncAAs, which is costly, not always suitable for large scale production, and is not environmentally friendly^42,43^. We reasoned that our evolved RebH_Evo4_ could tackle all of these problems by allowing both screening for active variants and their subsequent large-scale production to be performed autonomously in *E. coli*.

First, we constructed an arabinose-inducible peptide expression system which, in concert with Fre-L3-RebH_Evo4_ and the ChPheRS4 synthetase, allowed us to express peptides containing arbitrary, site-specific 7-Cl-Trp or 7-Br-Trp modifications, thereby enabling a suicide assay for AMP variants that impair growth rate (Figure 5C). Taking the recently reported AMP Enterocin RJ-11 as a model^44^, we rapidly screened for growth-inhibiting activity with chlorination at each of its 3 tryptophan residues (W12, 30, and 38) and observed that modification at position 30 resulted in no significant loss of potency (Figure 5D).

To confirm these results, we performed a minimum inhibitory concentration (MIC) assay on commercially-synthesised versions of WT Enterocin RJ-11 and its Trp30-chlorinated counterpart, and saw no significant difference between their ability to kill *E. coli* cells, indicating that the results of our screening approach can accurately predict halogenated AMP activity. Using a previously-reported SUMO-fusion approach^45^, we then overexpressed, cleaved, and purified our Trp30-chlorinated variant, confirming via HPLC and mass spectrometry that the product was the correct, chlorinated species. The biosynthesised peptide killed cells effectively via the same mode of action (membrane damage) as a commercially chemically-synthesised control (Figure 5H). Together, these results demonstrate that our evolved halogenase can enable efficient production of halogenated proteins and peptides, without the need to supplement the media with expensive ncAAs.

## Discussion

Engineered halogenases have long promised the possibility of bioproducing a wide variety of halogenated products, but attempts to overcome the inherent limitations imposed by their poor solubility, temperature stability, and catalysis have been difficult. Using phage-assisted continuous evolution and our GFP biosensor and biosynthesis circuits, we successfully engineered a RebH variant which to our knowledge shows the highest activity *in vivo* of any tryptophan halogenase currently reported. Our final evolved halogenase, RebH_Evo4_, has 12 mutations distributed across its surface, internal structure, and around the flavin binding pocket which, in addition to increasing its activity, make RebH_Evo4_ significantly more soluble than its WT counterpart — to our knowledge, this is also the first reported RebH variant with increased solubility.

Previous efforts to improve halogenases through directed evolution largely employed low-throughput screening of *in vitro* enzyme activity^20,46,47^, where mutations may arise that do not translate well to in *vivo* use^48^. This is particularly relevant in the case of RebH, which as a metabolically-coupled enzyme requires multiple other purified enzymes and cofactors to function *in vitro*, making *in vivo* use more practical for large-scale, low-cost bioproduction^12,39^. By employing an *in vivo*, phage-coupled selection using a biosensor, we not only improved the amount of sequence space explored in evolution by multiple orders of magnitude, but also ensured that the mutations we identified would confer real-terms activity increases in the intended system of use.

Since halogenation is often the bottlenecking step in biosynthetic pathways^7,13^, our demonstration that RebH_Evo4_ is over an order of magnitude (∼40x) improved in both halogenated tryptophan and tryptamine biosynthesis represents an important step towards the sustainable bioproduction of multiple highly relevant halogenated drugs^35^, pigments^7^, and industrially-useful molecules^11,8,13,12^. Furthermore, halogenated tryptophan is uniquely also able to be incorporated into peptides and proteins as a ncAA, where it can serve as a stability and activity enhancing modification. Recent advances in genetic code expansion enable efficient incorporation of multiple ncAAs into regular and macrocyclic peptides, but still require their direct supplementation to growth media, which is often cost-prohibitive^49–51^. Here, we demonstrate that our evolved RebH_Evo4_ facilitates both in vivo screening and autonomous bioproduction of potent halogenated antimicrobial peptides without the need for supplementation with expensive ncAAs, paving the way for inexpensive development and biomanufacturing of next-generation peptide antibiotics.

The success of our approach may be valuable for engineering of enzymes more broadly. Most biosensors employed to date for selection and evolution of improved enzyme variants have been transcription factors or riboswitches^52–55^, whereas the use of aaRSs as biosensors has remained relatively underexplored. Interestingly, hundreds of different ncAAs have been selectively incorporated into proteins in a highly specific manner using evolved aaRS-tRNA pairs^49,56^, which represents a significant untapped opportunity to evolve and engineer a large number of pathways for the biosynthesis of valuable ncAAs or their derivatives. Moreover, unlike transcription factor-based biosensors, aaRS-based readouts enzymatically consume the product they are detecting, potentially increasing their ability to select for highly active enzyme variants by better protecting against diffusion between neighboring cells. aaRSs are uniquely well suited to distinguish between similar small molecules, making them a promising avenue for further sensor engineering where precise delineation between multiple possible enzyme products is essential.

We envision our evolved enzyme’s improvements in solubility and activity serving as a chassis for next-generation halogenase design. Notably, none of the twelve adaptive mutations in RebH_Evo4_ fall within its tryptophan binding pocket; an interesting future line of work would combine this with previously reported efforts in engineering RebH to alter its substrate scope. One can imagine combinatorial engineering campaigns marrying the high *in vivo* activity and solubility of RebH_Evo4_ with machine-learning guided specificity mutations or other aaRS-based biosensors. The remarkable improvements of RebH_Evo4_ remove a key obstacle in the path towards scalable bioproduction of halogenated therapeutics, materials and bioactive peptides.

## Materials and methods

### Bacterial growth conditions, plasmid and phage cloning

For regular growth, cloning, and bacteria propagation, LB media was routinely used, and the correct antibiotics matching plasmids resistances were added to plates and media. Whenever more than 3 antibiotics were used in combination, antibiotic concentration was used as 0.5x of the final concentration. For K-12 Δ*tnaA* cells + F plasmid creation, K-12 Δ*tnaA::nptII*(KanR) cells (Keio collection) were conjugated with S2060 strain (which contains the F-plasmid) and selected against Kanamycin and Tetracyclin (marker present in S2060 F-plasmid).

Plasmids were cloned using NEBuilder HiFi assembly (New England Biolabs) and PCRs were performed using Q5 2x Hotstart mastermix (New England Biolabs). For phage cloning, the phage backbone was amplified using PCR and gene fragments were inserted using NEBuilder. The transformation mix was added to the chemocompetent S238 strain, transformed by heat shock and then cells were grown overnight at 37°C in LB media without any addition of antibiotics. Next day, the culture was spun down and the supernatant filtered through a 0.22 µm syringe filter. The supernatant was then plaqued using previously described methods^57^, using Blue-o-Gal and incubated overnight at 37°C to grow. Individual plaques were then sequenced and grown with S238 strain for propagation, sequence-verified phage supernatant kept at 4°C.

For phage library preparation, a similar method described by Jones and collaborators^58^ was used. Briefly, error-prone PCR of RebH was performed using Agilent Mutazyme II kit, and cloned into our phage backbone using NEBuilder. A 100 µL final reaction volume was used for that, aiming to have a 1µg total of DNA in the reaction. After incubating for 30 min at 50°C, the assembly reaction was purified using Monarch PCR purification kit (NEB) and eluted in 10 µL of water. For competent cell preparation, 50mL of S238 cells were grown in 2xYT media until OD of 0.5. Cultures were then chilled for 10 min in ice and washed twice with ice-cold 10% glycerol, and resuspended into a final volume of 100 µL. Cells were then mixed with the whole purified assembly reaction and transformed via electroporation, and recovered immediately in SOC media. Estimation of the number of cloned phage was performed after 30 to 45 min of recovery. Cells were left to grow overnight and next day, the phage library suspension was collected and treated as described above.

### Amber suppression sfGFP assays

Amber suppression was performed using either K-12 Δ*tnaA* strains or DH10B (New England biolabs) cells. For amber suppression experiments, a mixture of M9 media (M9 + casamino acids (2% w/v final) + glucose (0.5% w/v) and Davis Rich Media (DRM)^59^ was used, in a proportion of 9:1. Cells were pre-grown into M9/DRM mixture to exponential phage and then inoculated into fresh media, supplemented with 0.5 mM IPTG (final concentration) and the corresponding antibiotics. Cells were then grown overnight, spun down and washed twice (8000xg, 3min) with PBS, before reading the sfGFP signal using a TECAN plate reader, using a 96-well plate.

### Phage propagation assays and quantification

Phage propagation was estimated either by plaquing or by using qPCR to estimate phage enrichment. For plaquing, the same method described above was used, and previously described^57^, always as an overnight incubation at 37°C. For qPCR phage estimation, 0.75μL of phage suspension was combined with 5μL of PowerUp™ SYBR™ Green Master Mix (Thermo Fisher Scientific), 0.0625 μl each of 100 μM M13 forward and reverse primers (5′-CACCGTTCATCTGTCCTCTTT-3′ and 5′-CGACCTGCTCCATGTTACTTAG-3′), and water to achieve a final volume of 10 μL. The qPCR was performed with the following conditions: 95 °C for 2 min, and then 40 cycles of 95 °C for 15 s and 60 °C for 60 s. To generate a standard curve for qPCR, a standard phage sample of high titre (∼1 × 10^10^ plaque-forming unit (p.f.u.) per mL as determined by plaquing) was serially diluted by a factor of 10, up to 10^7^-fold, in water. A standard curve was generated using Cq values and phage titres were determined accordingly.

### Phage-assisted continuous evolution (PACE)

PACE was performed using similar methods as described previously^57^, with a few alterations. In our PACE system, instead of using a chemostat providing new host cells to the lagoon, we used a cell culture kept at 4°C. Briefly, cells that were freshly transformed with the MP6 mutagenesis plasmid^31^ were grown in M9:DRM media until reaching OD 0.05 to 0.1, then quickly chilled on ice and subsequently placed at 4°C. We observed that bacteria cells would grow and divide inside the tubing, especially in the section outside of the fridge and in the section inside the 37 degrees incubator, before reaching the lagoon, giving us a higher final OD (around 0.25 to 0.5) inside the lagoon. Culture media with inducer (L-arabinose) was kept in a separate flask and mixed directly into the lagoon via separate tubings, to achieve induction of the mutagenic proteins encoded on MP6 inside the lagoon. Peristaltic pumps were used to control the flow of culture and also media removal from the lagoon.

### RebH solubility assays

To test RebH solubility, we cloned the WT sequence and our evolved variant into a pET28a vector (pAP108 and pAP109, Supp. Table 1). The His-Tag sequence was removed. We then transformed BL21(DE3) cells, and grew independent colonies in 2xYT until reaching OD 0.4. Protein expression was then induced for 3 hours by adding IPTG (1 mM) to the media. After induction, cell pellets were harvested for protein extraction. Total protein fraction and soluble fraction were prepared by using the exact same volume of culture, similarly as reported previously^7^. For total protein fraction, cells were pelleted and resuspended in a 1x Laemmli buffer (BioRad). For soluble protein fraction, cells were pelleted, frozen and thawed and resuspended in B-PER protein extraction reagent (Thermo Scientific). Extraction with B-PER was performed at RT for 1 hour at a rotary well. The extract was then centrifuged (16,000xg for 5 minutes) and the supernatant was collected. Samples were loaded onto an SDS-PAGE gel (4-18%), run using MOPS buffer and stained using Coomassie blue.

### HPLC/MS quantification of 7-halotryptophan and 7-halotryptamine bioproduction

For HPLC/MS estimation and quantification of 7-halotryptophan and 7-halotryptamine production, *E. coli* cells (K-12 Δ*tnaA*) cells were freshly transformed with the respective plasmids containing the WT RebH and RebH_Evo4_ sequences. Both systems used the same genetic arrangement in an operon, with J23100 as a promoter and RBSc44 for Fre and SD8 for RebH variants. (See plasmid pAP65 and pAP113, Supp. Table 1 and 2). For 7-halotryptamine assays plasmid pJB21×05 encoding RgnTDC was additionally co-transformed. Colonies from each plate were collected and inoculated into LB media either at 37°C or at 30°C. Cells were then grown to an OD of 0.55 (±0.05), pelleted and resuspended to an OD of 5 in the final M9-based whole cell catalysis buffer. Base Cl-free M9-derivative (Buffer v4.1): Na₂HPO₄ (47.8 mM), KH₂PO₄ (22.0 mM), (NH₄)₂SO₄ (9.4 mM), casamino acids (0.2% w/v), glycerol (0.4% v/v), glucose (25 mM). Additionally, 5 mM tryptophan substrate was used in all assays, and 200 mM NaCl or NaBr was used depending on the intended product. The final volume of the reaction was 2mL. Cells were incubated in 50mL falcon tubes, overnight at 280 RPM either at 37°C or 30°C. We observed that strong aeration correlates with good production of halogenated products.

Samples were centrifuged for 10 minutes at 4000 xg and 30 µL of the supernatant were taken for HPLC/MS analysis, LC-MS data were obtained on a Waters ACQUITY (Massachusetts, USA) equipped with QSM, QDa and PDA detectors, sample manager FTN-H, quaternary solvent manager, column manager with ACQUITY UPLC BEH C18 1.7 μm, 2.1 x 50 mm column. Electrospray ionisation (ES+ and ES-) and Diode Array spectra were obtained for each characterised compound. The gradient methods for LC-MS were composed of 0.1% formic acid (FA) in water/ 0.1 % FA in acetonitrile, over 8 minutes. Peak identity was verified by Mass Spectrometry and quantification of the peaks (280 nm) was performed by comparing peak area to a standard curve obtained by running commercial standards of 7-Cl-Trp and 7-Br-Trp. Experiments were performed in triplicates.

### sfGFP purification and MS analysis

For sfGFP purification and MS, *E. coli* cells (K-12 Δ*tnaA*) transformed with plasmid containing RebH_Evo4_ and ChPheRS-4*S333C+M490L were grown into LB media until OD 0.1. Cells were spun down, washed 2x in PBS and resuspended in a 20mM sodium phosphate buffer, with either 200mM NaCl or 200mM NaBr, 1% glucose and 2g/L of casamino acids. IPTG was added to a final concentration of 0.5mM. Cells were grown overnight at 37°C. Next day, the pellet was collected and GFP was extracted using B-PER (Thermo Scientific) and purified using HisPur Ni-NTA resin (Thermo scientific). The purity of the purified sfGFP was analyzed using a SDS-PAGE gel, followed by staining with coomassie-blue. The sfGFP was then analyzed by MS and the intact mass analyzed.

### Fed-batch bioreactor production of 7-Cl-Trp

Scale-up to 5L bioreactor cultivation closely mirrored initial overnight shake-flask development, with a few modifications. The IPTG-inducible high-expression promoter proK-lacO was cloned upstream of the Fre, RebH operon to mitigate toxicity associated with constitutive expression. To initiate the bioreactor runs a pre-culture of freshly-transformed cells was inoculated in 24L in a 100L Eppendorf Bioflo 610 bioreactor, with appropriate antibiotics and 25 mM glucose in Terrific Broth until OD 0.50, whereupon cells were induced with 0.5 mM IPTG and allowed to grow again until they reached OD 1.00. 24L of the resulting pre-culture was spun down at 7340 ×g for 30 minutes at 25°C (2L Sorvall RC 12BP+ centrifuge) and resuspended to an OD of 4.8 in 5L of our M9-based transformation buffer v4.1 (described above) and transferred to the final bioreactor (10L Sartorius Biostat C bioreactor). Additional components used for 7-Cl-Trp fermentation: tryptophan (20 mM), IPTG (0.5 mM), NaCl (200 mM). 1 mL samples were taken manually every hour and OD recorded. Feed consisted of glucose (500 g/L), (NH_4_)_2_SO_4_ (100 g/L), and K_2_HPO_4_ (5 g/L), and dosed via automatic feedback control via rDOS probe, with a programmed setpoint of 25%. Samples were taken manually every hour. Since we had previously noticed that we had been reaching the limits of 7-Cl-Trp solubility at the titres observed by the end of the run, pellets were separated, extracted using acetonitrile, and unified with supernatant to account for all product that had precipitated prior to HPLC/MS quantification (as described above).

### Growth-based screening of halogenated antimicrobial peptide variants

Plasmids containing halogenation machinery plus the CDS of AMP variants with Trp residues selected for halogenation replaced with TAGs were cloned, with peptide CDSs under the araBAD promoter and bearing an SD8 RBS (see pJB15×03-series in Supp. Table 1). Colonies of NEB 10-beta cells freshly cotransformed with these plasmids plus the synthetase machinery plasmid described above (see pAP80 in Supp. Table 1) were grown until OD 0.50 in M9 + Casamino acids (0.2% w/v) and Glycerol (0.4% v/v), supplemented with 25 mM Glucose and appropriate antibiotics, then diluted 100X into 150 µL of the same base medium +/-0.5 mM IPTG and 10 mM Arabinose in 96-well optical flat bottom reader microplates (Corning). Growth curves were measured over 10hrs of continuous shaking at 37°C on a Tecan Spark microplate reader.

### Autonomous expression and purification of halogenated Enterocin RJ-11

Plasmids bearing halogenation machinery plus the CDS for Trp30-chlorinated Enterocin RJ-11 fused C-terminally to a 6xHis-SUMO tag were cloned, with the SUMO-peptide CDS under the T7 promoter and bearing an SD8 RBS (see pJB18×02v1 in Supp. Table 1). C321.ΔA T7RNAP ompT-rne-lon-cells (Addgene #182778^60^) were cotransformed with this plasmid plus the synthetase machinery plasmid described above (see pAP80 in Supp. Table 1), and were grown until OD 0.50 in M9 +

Casamino acids (0.2% w/v) and Glycerol (0.4% v/v), supplemented with 25 mM Glucose and appropriate antibiotics. SUMO-peptide expression was induced overnight with 0.5 mM IPTG, and pellet harvested, resuspended using B-PER Complete Lysis Buffer (Thermo), and sonicated. SUMO-peptide was purified from clarified lysate using Ni-NTA magnetic beads (NEB), cleaved with ULP-1 (Sigma) for 1hr at 30°C, and then reverse-purified using a second round of IMAC. The final purified peptide was buffer-exchanged into deionised water using a 3MWCO centrifugal filter (Sartorius). The samples were analysed by HPLC/MS on an Agilent 1260 LC-MSD (Agilent Poroshell 120 EC-C18 2.7um, 3.0×30mm column), using a linear gradient of 5-100% B over 8.5 min at a flow rate of 0.425 mL/min. A binary solvent system [A: and H2O /0.08% TFA/1% MeCN and B: MeCN /0.08% TFA] was used.

### MIC assays for antimicrobial peptides

*E. coli* cells (NEB 10-beta) were grown to OD 0.50 in LB without antibiotics, then diluted 100X into 300 µL LB in round bottom 96-well deep culture plates, supplemented with a dilution series of the appropriate peptide. Commercially-synthesised peptides (Biosynth) were prepared at 128 µM in deionised water. Plates were incubated overnight at 37°C with 280 RPM shaking covered with a Breathe Easier (Sigma) membrane, then 100 µL of each was transferred to a 96-well optical flat bottom reader microplate (Corning) and OD_600_ measured on a Tecan Spark microplate reader. Measurements were normalised as a percent relative to growth and sterility controls included in every assay.

### Fluorescence microscopy membrane integrity assays

For fluorescence microscopy observations, *E. coli* cells (K-12) were grown in LB media until exponential phase (OD 0.5). Cells were then treated with either the commercial Enterocin RJ-11_W307-Cl_ or the biosynthetized and purified Enterocin J-11(W30) 2x MIC value (∼3 µM) and incubated for 15min at 37°C with shaking. Cells were then harvested (1mL), centrifuged for 3min at 4000xg and resuspended in 100ul of residual media. Cells were treated with 50 µg.ml−1 of FM1-43 (Invitrogen^TM^) and 10 µg. ml−1 of propidium iodide for membrane staining and permeability evaluation, respectively. About 5µl to 10µl of cell suspension were added on top of LB 25% agarose pads, covered with coverslips and imaged using a Nikon Ti2 Eclipse inverted microscope, with a 100x oil objective. Images were analysed using ImageJ.

## Supporting information

Plasmid Maps in .gb

## Author Contributions

A.A.P. and E. D. conceived the study. A.A.P, J.B. and E.D. designed the experiments A.A.P., J.B., and L. N. cloned the plasmids. A.A.P. created and calibrated the microbial circuits, performed phage and evolution experiments. O.A. and S.A. established phage methods and PACE set-up. A.A.P., J.B., A.S. and C.S., performed HPLC/MS measurements. J.B. and J.C. performed protein/peptide purification and MS. J.B., N.P. and A.A. performed bioreactor experiments. J.B. performed antimicrobial peptide screening and purification. A.A.P. and J.B. performed microscopy. Figures were prepared by A.A.P. and J.B. with input from all other authors. Text and were prepared by A.A.P. , J. B. and E. D. with input from all other authors.

## Supplemental Information

**Supp. Table 1.** Plasmids used in this study. Plasmids used in this study, and the respective panels in which they were used. Full plasmid maps for every construct are available in the supplemental data file. Selected plasmids (*) are available on Addgene under accession numbers 247093 – 247103

| Figure | Plasmids used |
| --- | --- |
| Figure 1B, 1C and 1D | pAP79* + pAP49 (Biosensor circuit) |
| Figure 1F | pAP79* + pAP49 (Biosensor circuit)<br>pAP79* + pAP65* (RebH WT + Fre)<br>pAP79* + pAP62 (MBP + Fre) |
| Figure 2B, 2C | pAP79* (AP1) + pAP70* (AP2)<br>pAP77 (empty phage genome)<br>pAP68 (RebH phage genome) |
| Figure 3B | pAP79* (AP1) + pAP70* (AP2) + MP6<br>pAP68 (RebH phage genome)<br>pAP93 (RebH<sub>Evo1</sub> phage genome)<br>pAP105 (RebH<sub>Evo3</sub> phage genome) |
| Figure 3D | pAP79* (Biosensor)<br>pAP65* (RebH WT) + pAP79*<br>pAP94 (RebH<sub>Evo1</sub>) + pAP79*<br>pAP102 (RebH<sub>Evo2</sub>) + pAP79*<br>pAP104 (RebH<sub>Evo3</sub>) + pAP79*<br>pAP113* (RebH<sub>Evo4</sub>) + pAP79* |
| Figure 4A, 4B | pAP65* (RebH WT)<br>pAP113* (RebH evo4) |
| Figure 4C | pAP108 (RebH WT under T7 promoter)<br>pAP109 (RebH<sub>Evo4</sub> under T7 promoter) |
| Figure 4D | pAP65* (RebH WT) + pAP79*<br>pAP54* (Fre-L3-RebH WT) + pAP79*<br>pAP113* (RebH<sub>Evo4</sub>) + pAP79*<br>pAP114* (Fre-L3-RebH<sub>Evo4</sub>) + pAP79* |
| Figure 4E | pAP114* (Fre-L3-RebH<sub>Evo4</sub> GFP*1TAG) + pAP80* |
| Figure 5A | pAP65* (RebH WT) or pAP113* (RebH<sub>Evo4</sub>)<br>pJB21x05* (RgnTDC) |
| Figure 5B | pJB23x02 (proK-lacO RebH<sub>Evo4</sub>, Fre) |
| Figure 5C, 5D | pJB15x02* (Enterocin RJ-11 WT)<br>pJB15x06v1, 2*, 3 (Enterocin RJ-11 W12, 30, 38 TAG) |
| Figure 5F, 5G | pJB18x02v1* (T7-lacO SUMO-Enterocin RJ-11 W30 TAG)<br>pAP80* |
| Supp. Figure 1A | pAP58 (ChPhe tRNA) + pAP49<br>pAP63 (tRNA<sub>3C11</sub>) + pAP49 |
| Supp. Figure 1B | pAP63 (tRNA<sub>3C11</sub>) + pAP49<br>pAP79* (*S333C + tRNA<sub>3C11</sub> tRNA) |
| Supp. Figure 2B, 2D, 2E | pAP65* (RebH WT) + pAP79*<br>pAP94 (RebH<sub>Evo1</sub>) + pAP79*<br>pAP102 (RebH<sub>Evo2</sub>) + pAP79*<br>pAP104 (RebH<sub>Evo3</sub>) + pAP79*<br>pAP113* (RebH<sub>Evo4</sub>) + pAP79* |
| Supp. Figure 3 and Supp. Figure 4 | pAP65* (RebH WT)<br>pAP113* (RebH<sub>Evo4</sub>) |
| Supp. Figure 5A | pAP118 (WT GFP)<br>pAP54* (Fre-L3-RebH WT) + pAP80*<br>pAP114* (Fre-L3-RebH<sub>Evo4</sub> GFP*1TAG) + pAP80*<br>pAP119 (Fre-L3-RebH<sub>Evo4</sub> GFP*3TAG) + pAP80* |
| Supp. Figure 5B | pAP114* (Fre-L3-RebH<sub>Evo4</sub> GFP*1TAG) + pAP80* |
| Supp. Figure 6A | pJB15x02* (Enterocin RJ-11 WT)<br>pJB15x06v1, 2*, 3 (Enterocin RJ-11 W12, 30, 38 TAG) |
| Supp. Figure 6B | pJB18x02v1* (T7-lacO SUMO-Enterocin RJ-11 W30 TAG)<br>pAP80* |

## Supplemental Figures

**Supp. Figure 1.**
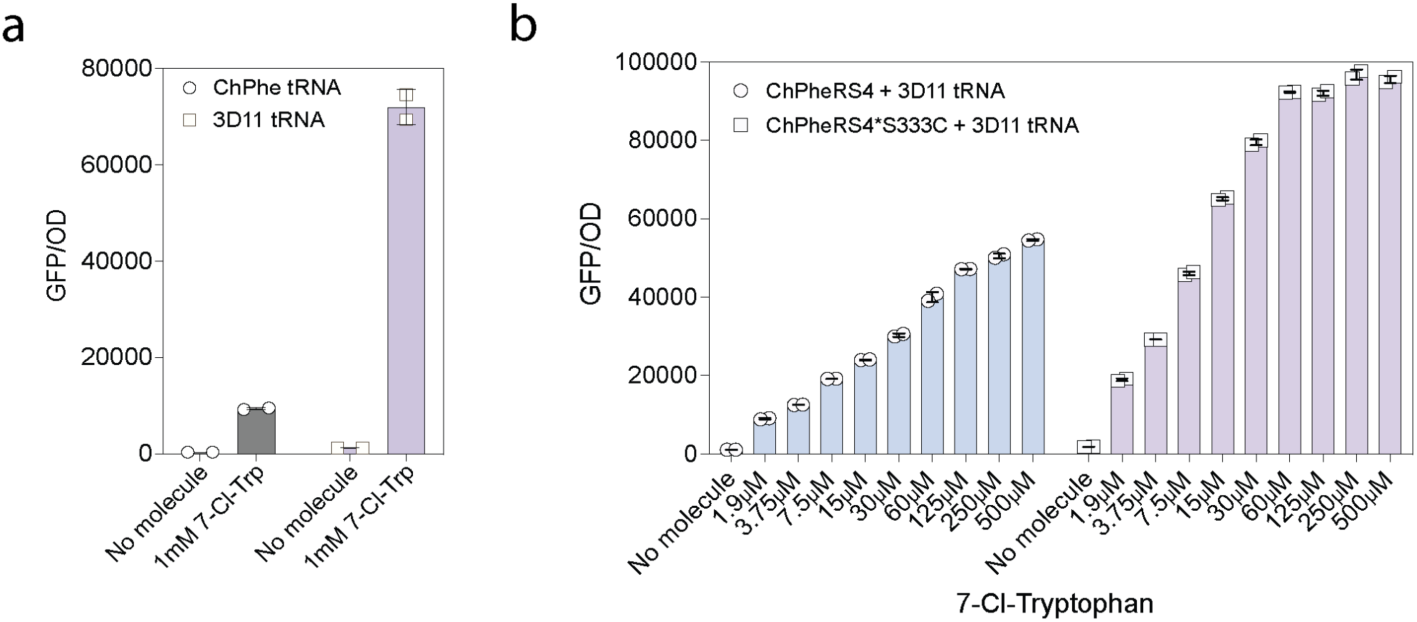
- SI 1a. Amber suppression of sfGFP using either the original ChPheRS4 tRNA, or the improved 3D11 tRNA, in the presence of absence of 7-Cl-Trp. - SI 1b. Amber suppression of sfGFP using either the original ChPheRS4, or an improved mutant ChPheRS4*S333C, in the presence or absence of 7-Cl-Trp.

**Supp. Figure 2.**
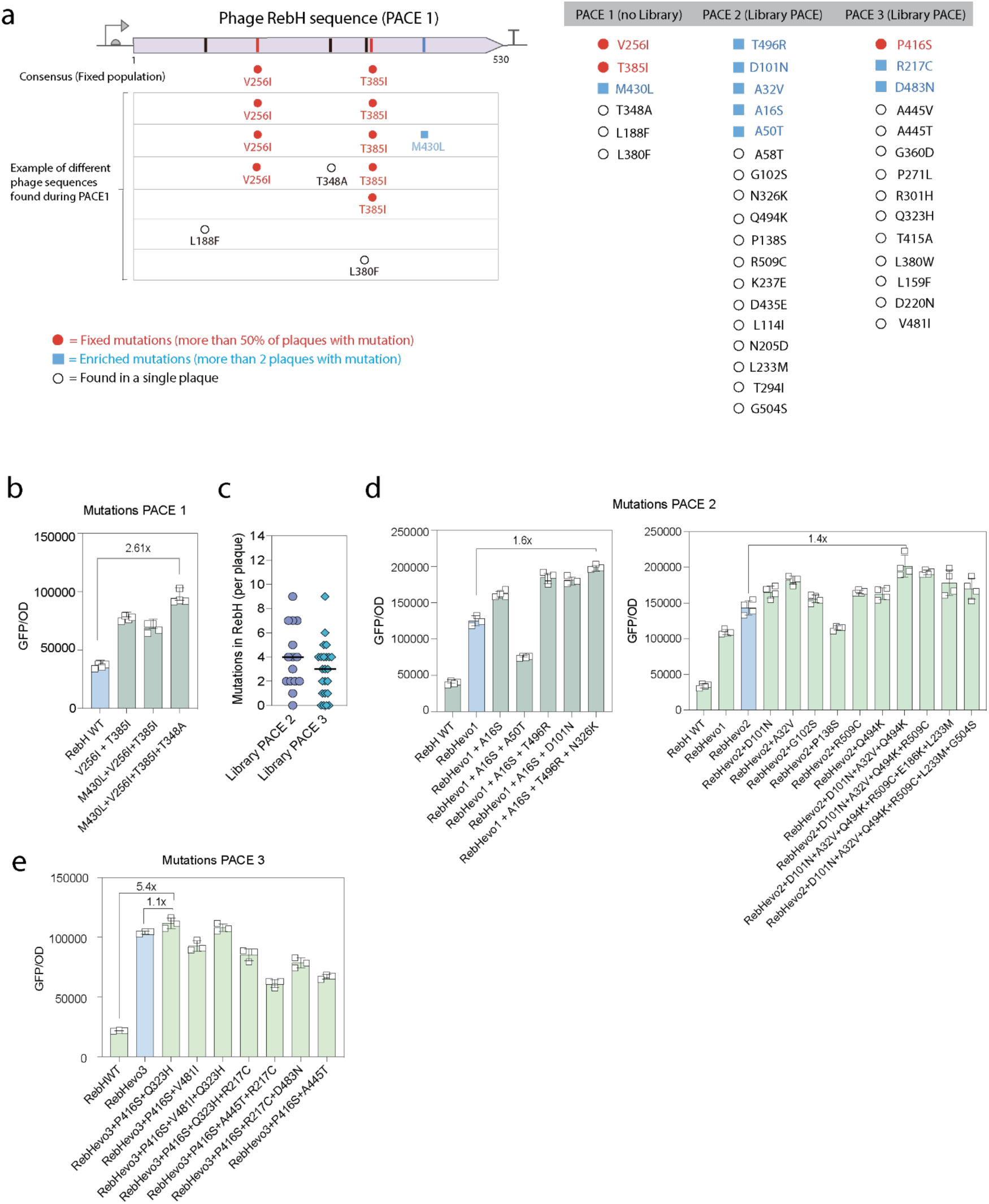
- SI 2a. Results from the first PACE run, highlighting the different mutations obtained, and their abundances (on the right). On the left, list of mutations recovered on clonal phage on each run. - SI 2b. Combination of different mutations obtained during PACE 1, using sfGFP circuit. - SI 2c. Estimation of number of mutations per plaque in each library used for PACE 2 and PACE 3, after sequencing individual plaques. - SI 2d. Combination of different mutations obtained during PACE 2, using sfGFP circuit. - SI 2e. Combination of different mutations obtained during PACE 3, using sfGFP circuit.

**Supp. Figure 3.**
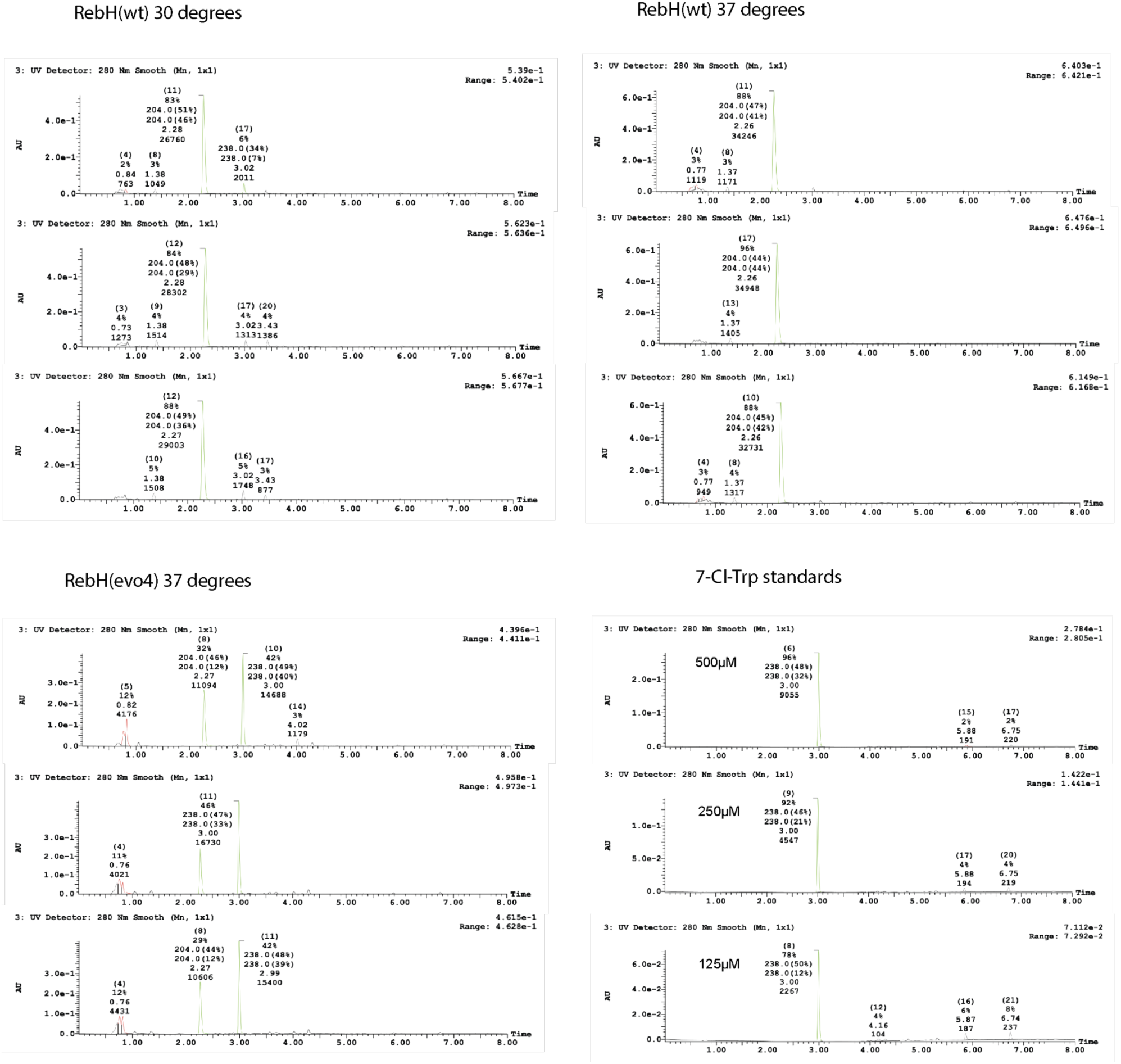
HPLC spectra from 7-Cl-Trp production using either RebH_WT_ or RebH_Evo4_

**Supp. Figure 4.**
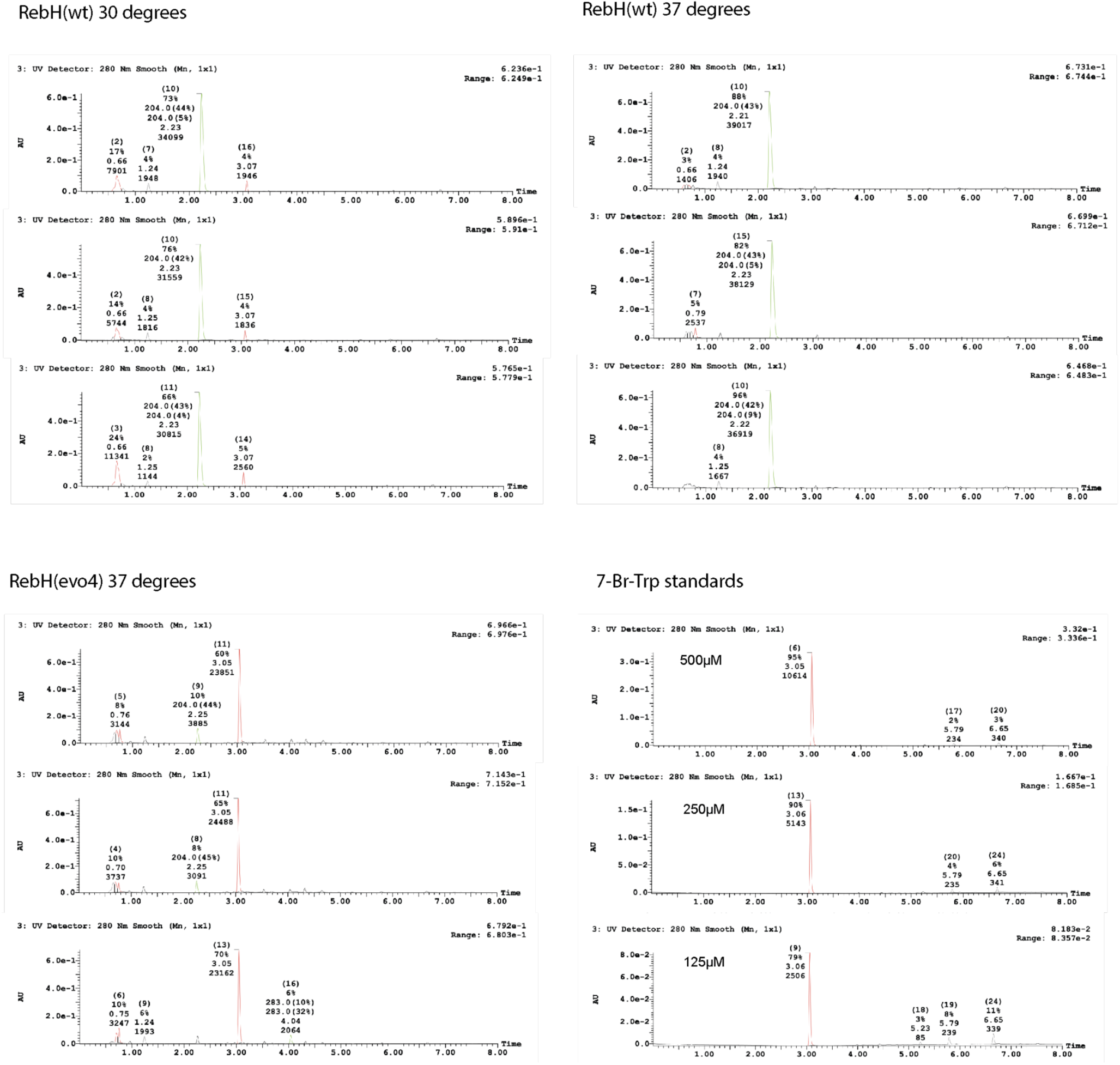
HPLC spectra from 7-Br-Trp production using either RebH_WT_ or RebH_Evo4_

**Supp. Figure 5.**
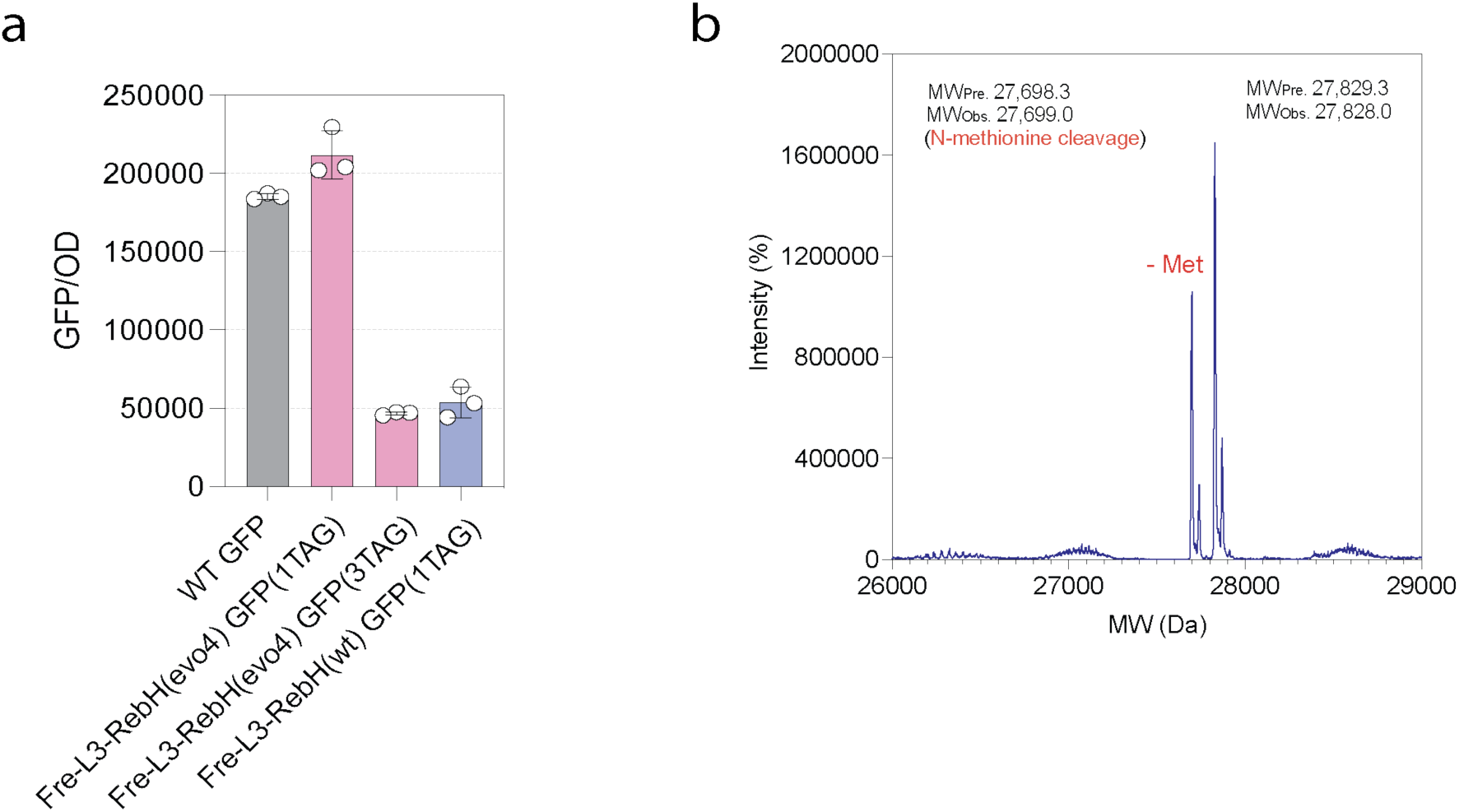
- SI 5a. Amber suppression of sfGFP using either Fre-L3-RebH_Evo4_ or Fre-L3-RebH_WT_. Amber suppression efficiency is much more efficient when using the evolved enzyme. - SI 5b. Intact Mass of sfGFP produced using Fre-L3-RebH_Evo4_ in media supplemented with NaBr. The observed mass matches the expected mass from sfGFP with 7-Br-Trp incorporated at position 151. MWpred. = Predicted molecular weight. MWobs: Observed molecular weight.

**Supp. Figure 6.**
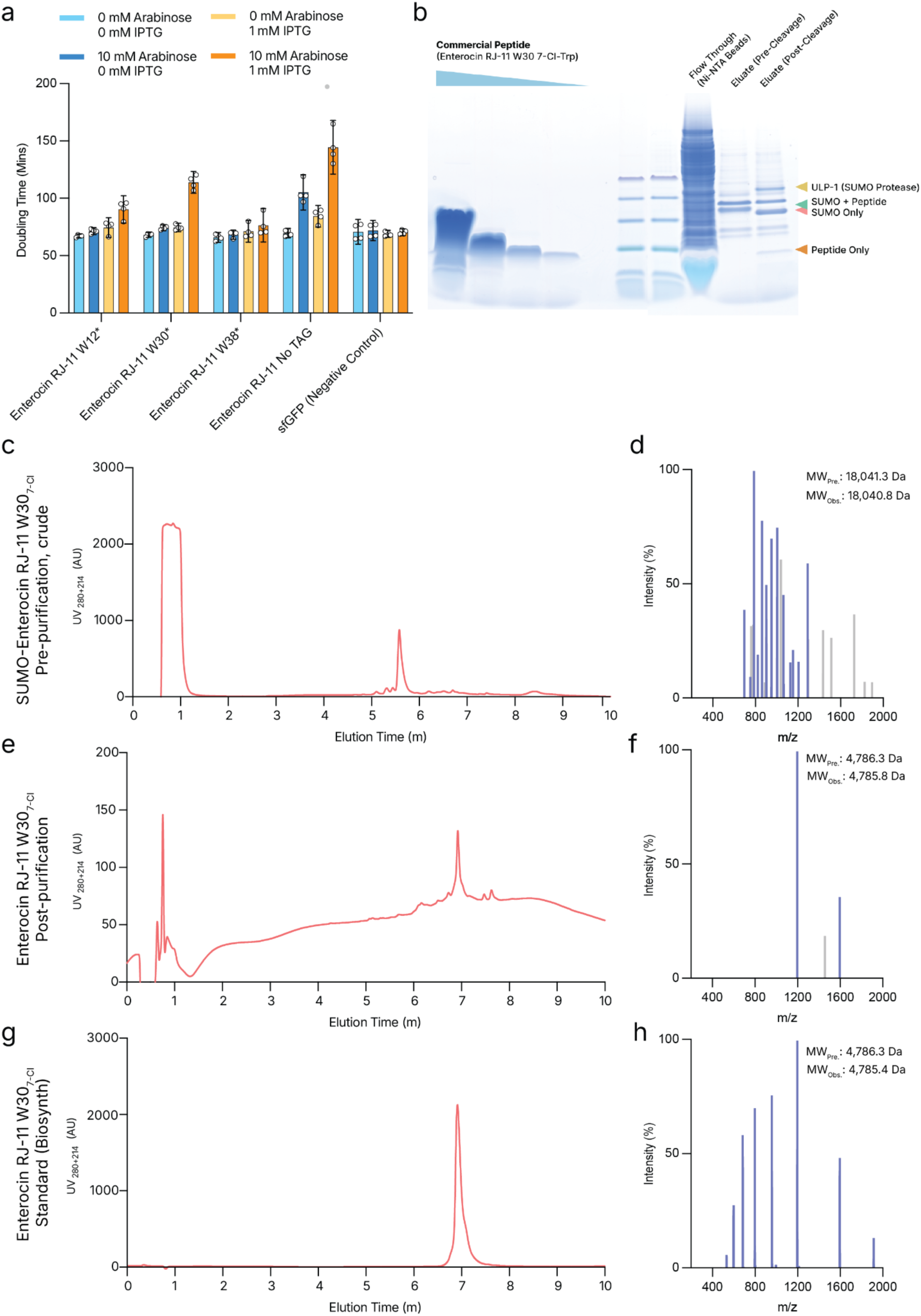
- SI 6a. AMP screening data from Figure 5D presented in bar chart as doubling time of AMP variant-expressing strains with and without AMP and ChPheRS4 expression (Arabinose and IPTG respectively) - SI 6b. Full SDS-PAGE gel from Figure 5F with commercially-synthesised peptide for comparison. - SI 6c,e,f. Full HPLC traces from Figure 5G showing crude SUMO-Enterocin RJ-11 W30_7-Cl_ pre-cleavage and purification, biosynthesised Enterocin RJ-11 W30_7-Cl_, and commercially synthesised peptide for comparison. - SI 6d,f,h. Mass-spectra used to calculate MW values presented in Figure 5G

